# Evolutionary dynamics of pseudoautosomal region 1 in humans and great apes

**DOI:** 10.1101/2021.09.14.460222

**Authors:** Juraj Bergman, Mikkel Heide Schierup

**Affiliations:** Bioinformatics Research Centre, Aarhus University, DK-8000 Aarhus C, Denmark

## Abstract

The pseudoautosomal region 1 (PAR1) is a 2.7 Mb telomeric region of human sex chromosomes. As the largest point of contact between the X and Y, PAR1 has a crucial role in ensuring proper segregation of sex chromosomes during male meiosis, exposing it to extreme recombination and associated mutational processes. We investigate PAR1 evolution using population genomic datasets of extant humans, eight populations of great apes and two archaic human genome sequences. We find that the PAR1 sequence is closer to nucleotide equilibrium than autosomal telomeric sequences. We detect a difference between long-term substitution patterns and extant diversity in PAR1 that is mainly driven by the conflict between strong mutation and recombination-associated fixation bias at CpG sites. Additionally, we detect excess C→G mutations in PAR1 of all great ape species, specific to the mutagenic effect of male recombination. Analysis of differences between frequencies of alleles segregating in females and males provided no evidence for sexually antagonistic selection in this region. Furthermore, despite recent evidence for Y chromosome introgression from humans into Neanderthals, we find that the Neanderthal PAR1 retained similarity to the Denisovan sequence, as is the case for the X chromosome and the autosomes. Lastly, we study repeat content and double-strand break hotspot regions in PAR1 and find that they may play roles in ensuring the obligate X-Y recombination event during male meiosis. Our study provides an unprecedented quantification of population genetic forces and insight into evolutionary processes governing PAR1 biology.

## Introduction

The mammalian sex chromosomes, X and Y, originated from a pair of autosomal precursors around 180 million years ago [1]. At least four subsequent recombination suppression events and loss of sequence homology in the sex-determining region (SDR) have resulted in extreme divergence of sequence and function between the sex chromosomes [2–6]. Yet regions with X-Y homology and genetic exchange, termed the pseudoautosomal regions (PARs), still persist across placental mammals. In great apes, the pseudoautosomal region 1 (PAR1) is just 2.7 Mb in length, but has the important role of ensuring proper meiotic segregation of X and Y. PAR1 is also one of the most evolutionarily dynamic regions of the genome. Obligate male crossovers are restricted to this physically small region during male meiosis, so recombination rates per base pair are extremely high, and the region shows high nucleotide diversity. Recent progress in genome assembly of the sex chromosomes and the availability of population genomic datasets has now made it possible to study divergence and diversity processes of this important region in detail.

The importance of pseudoautosomal regions is evident in the association of PAR1-specific mutations with various phenotypic consequences in humans. Since homologous pairing and exchange of genetic material is crucial for successful gametogenesis (at least one cross-over per chromosome pair is necessary for proper segregation), PAR1 is especially important for male fertility [7,8]. Additionally, polymorphisms in PAR1 are associated with various diseases, such as skeletal malformations due to variants in the *SHOX* gene [9,10], schizophrenia [11], bipolar disorder [12] and hematological malignancies [13]. Large genomic rearrangements in PAR1 have also been reported - examples include an approximately 300 kb deletion associated with acute lymphoblastic leukemia, spanning several genes [14,15], a 47.5 kb deletion in the enhancer region of the *SHOX* gene [16] and an extension of the PAR1 region on the Y chromosome through a translocation of a 110 kb region from the X chromosome [17].

Sequence evolution of the pseudoautosomal region after the split between the avian and mammalian lineages, and leading up to extant mammalian species, involved the formation of several evolutionary strata mediated by recombination suppression between the sex chromosomes [18–20]. Consequently, PAR1 of humans and great apes is a small genomic region evolving under a concentration of strong population genetic forces. Such a convergence of forces results in PAR1-specific patterns of mutations, nucleotide composition evolution and recombination that have been mostly studied for large human diversity datasets [21–23]. Here, we aim to provide a broader perspective on PAR1 evolution by including nucleotide divergence and diversity data from eight populations of great apes, as well as ancient hominid data. We quantify divergence rates, and deviations of sequence composition from nucleotide equilibrium, and infer substitution and diversity spectra. For comparison, we use autosomal telomeres, which are similar to PAR1 in having higher male recombination rates and GC content, typical of male-specific recombination processes [24,25].

A central focus of our study is the relationship between recombination and nucleotide diversity in PAR1. Recombination in PAR1 has been subject to several studies, including sperm-typing, pedigree-based and sequence-based approaches. The general conclusion is that recombination in PAR1 during spermatogenesis occurs at a rate approximately 17-fold higher than the genome-wide average (1.2 cM/Mb), while the difference between male and female rates in the region is around 10-fold [6,12,21]. Recombination rate differences can affect sequence diversity in several ways. Firstly, recombination mediates the effect of directional selection on linked diversity. A strong reduction of nucleotide diversity during selective sweeps is expected in low recombining regions [26,27], and similarly, reduction in diversity due to background selection is diminished with increasing recombination rate [28]. Additionally, recombination directly affects diversity due to its mutagenicity [22,29], which is likely an important factor of PAR1 evolution [30–32]. The type of nucleotide diversity caused by the recombination process may also differ from the diversity introduced into the genome through DNA replication errors [22]. Recombination is also associated with GC-biased gene conversion, whereby purine-pyrimidine mismatches that arise during meiotic chromosomal pairing are most often resolved into GC, rather than AT pairs [33–35]. This effect on nucleotide diversity will be strongest in highly recombining genome regions, and is indistinguishable from directional selection favouring GC content [36].

Polymorphisms in the pseudoautosomal regions may also be maintained due to balancing and/or sexually antagonistic (SA) selection. Theory predicts that the recombination rate with the SDR will determine the conditions under which SA polymorphisms can persist [6,37,38]. In general, a sexually antagonistic variant that is tightly linked to the SDR can be maintained in a population under a broader range of selection coefficients compared to a freely recombining locus. Furthermore, as the recombination rate in PAR1 decreases with the distance from the telomere, PAR1 loci that are potentially under SA selection are therefore more likely to be found closer to the pseudoautosomal boundary. Additionally, a peak of diversity is expected around a PAR locus under balancing or SA selection [39]. Potential SA loci have been identified in various animal species using population genomic data to calculate *F*_*st*_ values between females and males [40–44], and sexual antagonism specific to the PAR has been suggested for the plant *Silene latifolia* [45,46]. A recent study of sexual antagonism in human pseudoautosomal regions reached equivocal conclusions [23], and SA in PAR1 of other great apes has not been investigated.

Another point of interest is the following. Hybridization events between ancient humans resulted in peculiar patterns of sex chromosome evolution. On the one hand, the X chromosome is devoid of ancient DNA introgression tracks in modern humans [47,48], while the human Y chromosome invaded and replaced the neanderthal Y lineage [49]. We therefore studied PAR1 evolutionary patterns in ancient hominids, using high-coverage sequences of a Neanderthal and Denisovan individual [50,51].

Lastly, we study the factors that ensure the occurrence of a recombination event in pseudoautosomal regions. Studies in mice have shown that repeat elements play a pivotal role in determining the physical structure of the PAR during recombination [52–54]. We therefore did a comparative analysis of repeat elements between PAR1 and autosomal telomeres. Another important determinant of PAR recombination is the recombination motif-recognition protein PRDM9, which is known to facilitate meiotic double-strand breaks and operate in human pseudoautosomal regions [55], as well as to evolve rapidly between primate lineages [56–58]. We therefore explored sequence evolution of double-strand break hotspots in PAR1 to gain insight into the role of PRDM9 in PAR1 sequence evolution across great apes.

## Results

### PAR1 divergence is exceptionally high across the great ape phylogeny

We used the Progressive Cactus aligner [59] to produce an alignment between the human, chimpanzee, gorilla, orangutan and macaque PAR1 sequences. The macaque sequence was used as an outgroup to root the tree, and for estimating divergence rates within the great ape phylogeny since the branching of the great ape ancestor. The estimated phylogenetic tree and alignment are shown in Figure 1. For tree estimation, we excluded sites in coding and repetitive regions, as well as CpG islands and conserved sites across primates. This left us with an alignment of 239,799 putatively neutral sites across the great ape phylogeny. Since the branching of great ape ancestor into extant species, PAR1 appears to have evolved at different rates (Figure 1; Table 1). The divergence rate in the chimpanzee lineage is approximately 5% larger compared to the PAR1 sequence of humans and gorillas, while the orangutan PAR1 sequence is the least diverged with a 10-16% lower divergence compared to the other species. Since the human-chimpanzee split, the divergence along the chimpanzee lineage is 14% higher than the human rate. Nevertheless, PAR1 sequences evolve faster than autosomal ones; pairwise divergence estimates of autosomal sequences for the human-chimp (0.0137), human-gorilla (0.0175) and human-orangutan species pairs (0.034) [60] are on average 40% lower than our PAR1 estimates.

**Figure 1.**
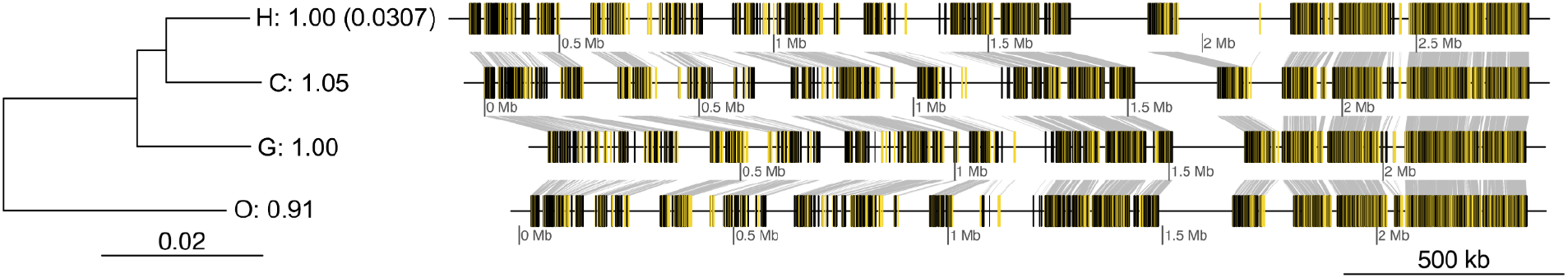
PAR1 phylogenetic tree and sequence alignment between great ape species. The total alignment consists of 620,054 syntenic sites of which 239,799 are putatively neutral (highlighted in yellow). As indicated by letters in the tree diagram, the species are as follows, from top to bottom: human (H), chimpanzee (C), gorilla (G) and orangutan (O). The number associated with each branch label is the divergence rate ratio with respect to the human rate (presented in parentheses), since the branching of the great ape ancestor.

**Table 1.**
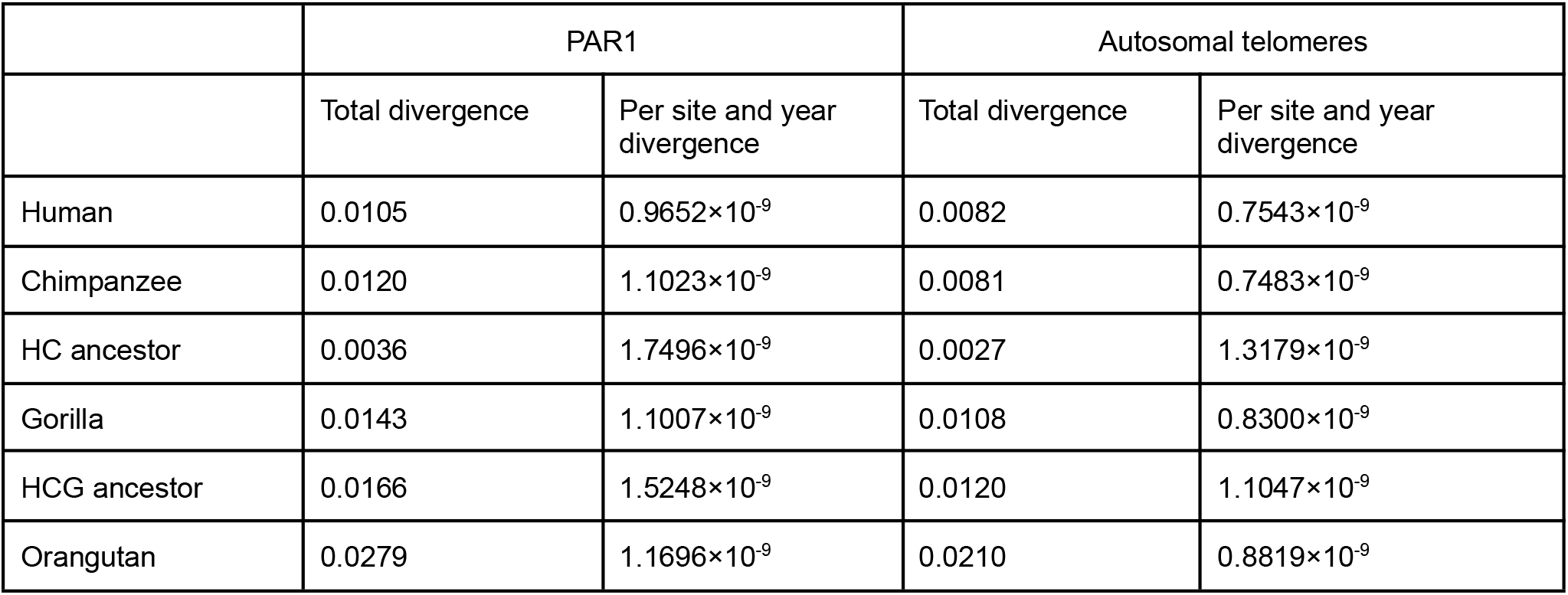
Divergence estimates based on phyloFit estimation and divergence times from [61] for the PAR1 sequence and the concatenated sequence of autosomal telomeres.

Based on estimated great ape divergence times [61], we estimate that the PAR1 divergence rate is within the range of 0.97-1.17×10^−9^ substitutions per site and year for the terminal branches of the phylogeny (Table 1). These rates are 1.51-1.83 fold higher than the yearly mutation rate for non-human great apes, estimated using trio sequencing (0.64 ×10^−9^ mutations per site and year; [61]). While it has been suggested that this mutation rate estimate is adequate for explaining autosomal divergence rates, the magnitude of PAR1 divergence implies additional mutational processes acting in this region. For comparison, we estimated the substitution rate for autosomal telomeric sequences (defined as 3 Mb syntenic regions that are present at the tips of autosomes in great ape species and the macaque outgroup). We concatenated sequences from all autosomal telomeres, in order to obtain a single estimate for each branch of the phylogeny. These sequences have a 1.16-1.38 fold higher divergence rate than the trio-based mutation rate, indicating that higher divergence is a general feature of telomeres, likely due to a combination of higher substitution rates and larger ancestral polymorphism caused by a higher local effective population size of telomeres. However, compared to autosomal telomeres, PAR1 has a 1.28-1.47 fold higher divergence rate across all branches of the phylogeny (Table 1), implying an exceptionally high rate compared to other fast-evolving regions.

### Nucleotide composition in PAR1 is closer to equilibrium than for other telomeres

We next consider PAR1 divergence as a function of derived state and substitution type. For comparison, and to obtain enough data, we computed divergence rates for each telomere separately. Specifically, we studied GC content evolution by comparing counts of AT→GC and GC→AT substitutions, and rates inferred from the posterior mean substitution counts between the corresponding nucleotides. These counts were inferred using the phyloFit program [62].

In line with the results in Table 1, the divergence rates of PAR1 in the four branches studied are higher than those for autosomal sequences, even when inferring divergence rates for each telomere separately (Figure 2). Although divergence rates of some autosomal telomeres overlap with PAR1 rates, the PAR1 rates are significantly higher (Wilcoxon *W* = 9, *p* = 0.0013). Generally, transitions occur more frequently than transversions, with the average ts:tv ratio of approximately 2.4:1 across all telomeres and species, which is somewhat higher than the autosomal estimate of 2.1:1 [63,64]. For PAR1 alone, the ts:tv is among the lowest of all telomeres (average of 1.92:1), ranging from 1.75:1 in the chimpanzee to 2.04:1 in the orangutan (Supplementary table 1).

**Figure 2.**
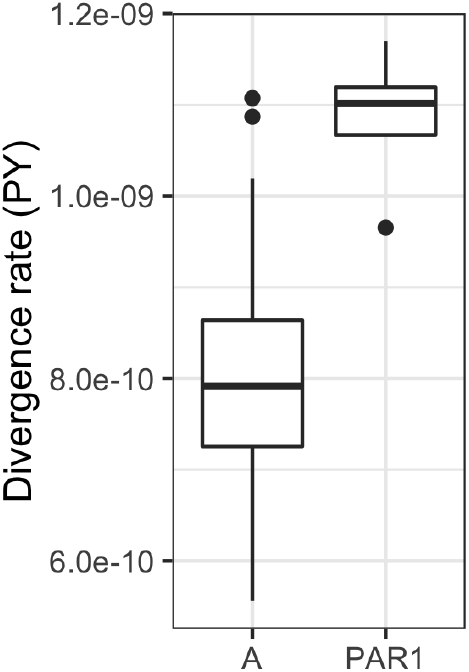
Distribution of per site and year (PY) divergence rates for autosomal (A) telomeres and PAR1 in the four branches studied, represented as box plots. For each telomere we included the four values corresponding to the four external branches of the great ape phylogeny. There were N=100 estimates for autosomes (across 25 alignable telomeres) and N=4 estimates for PAR1. Upper outliers for the autosomal estimates are the human and gorilla chromosome 8 telomeres, while the lower outlier for PAR1 is the human sequence.

To assess whether telomeric GC content is at equilibrium, we plotted the distribution of the ratio of AT→GC versus GC→AT substitution counts (Figure 3A). We expect this ratio to be close to 1 if the nucleotide composition of the sequence is close to equilibrium. However, it is below 1 for most telomeres, indicating that telomeric sequences are evolving towards lower GC content, as observed previously for autosomal sequences [65]. However, we observe that average AT→GC/GC→AT count ratios are greater in PAR1 compared to autosomal telomeres (Wilcoxon *W* = 317, *p* = 0.0245). We next calculate the equilibrium GC content of telomeres (GC*). The GC* value was calculated as the per site divergence rate for AT→GC mutations, divided by the sum of AT→GC and GC→AT rates, and it represents the stationary expectation for the proportion of GC nucleotides in a region evolving under the inferred divergence rates [65]. Figure 3B shows that the current PAR1 GC proportion is closer to its GC* value compared to the autosomal telomeres. PAR1 is likely closer to equilibrium both because its GC content has decreased more than for the other telomeres, and because its expected GC* is larger.

**Figure 3.**
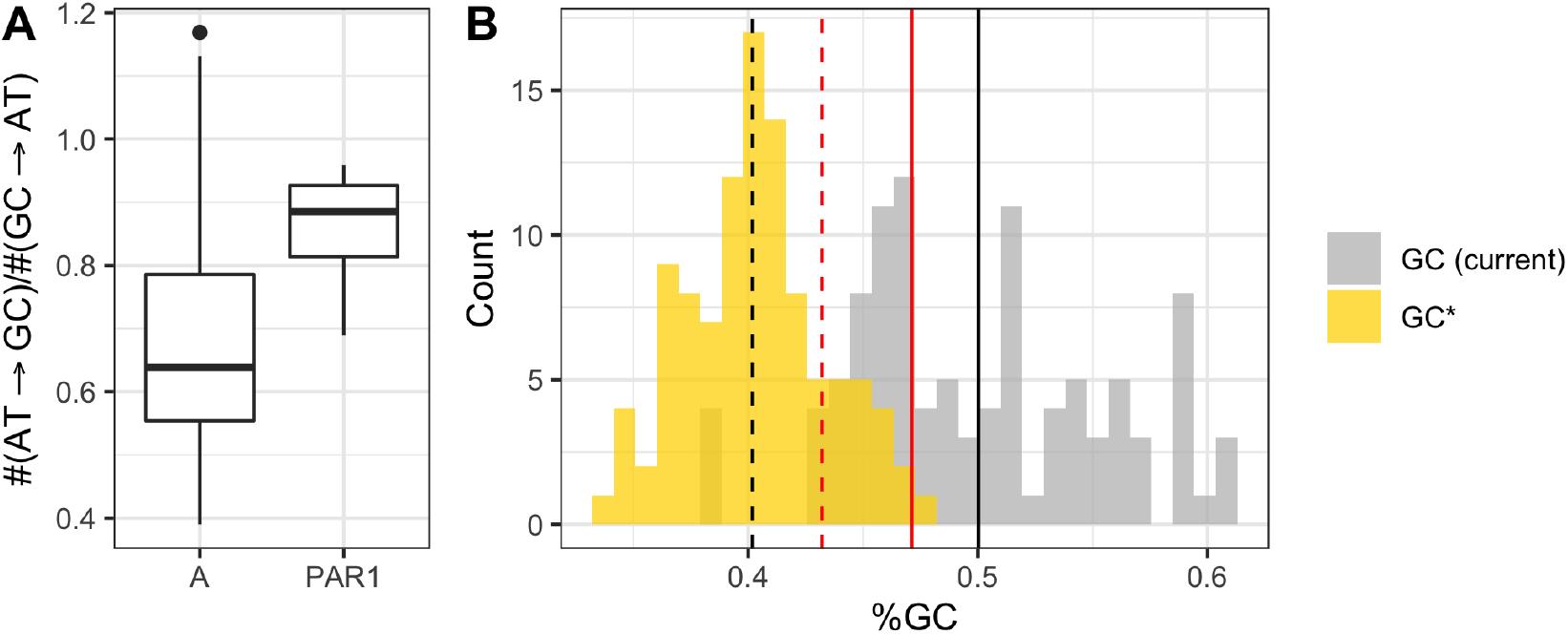
**A**. Distribution of AT→GC/GC→AT substitution count ratios for autosomal (A) telomeres and PAR1, represented as box plots. The upper outlier for the autosomal estimates is the gorilla chromosome 6 telomere. **B**. Distribution of current telomeric GC proportion and equilibrium GC proportion (GC*), expected given the inferred AT→GC and GC→AT substitution rates. The full and dashed black lines indicate the mean current and equilibrium GC proportion, respectively, for autosomal telomeres, and the red lines indicate analogous values for PAR1. For each telomere we plot four values corresponding to the four external branches of the great ape phylogeny. In total, we consider N=100 estimates for autosomes (across 25 alignable telomeres) and N=4 estimates for PAR1.

We next correlate the difference between GC* and current GC (*i*.*e*., a measure of deviation from nucleotide composition equilibrium) with divergence rate (Figure 4A), the ts:tv ratio (Figure 4B) and length of the chromosome upon which the telomere is located (Figure 4C). Partial correlation coefficients between these parameters are presented in Figure 4D. We observe that telomeres closer to equilibrium generally have lower divergence rates, with PAR1 as an obvious outlier. Similarly, the ts:tv ratio is negatively correlated to GC difference, with PAR1 having some of the lowest ts:tv values. On the other hand, chromosome length is positively correlated with GC difference. This is due to a larger current GC for small chromosomes and not differences in GC*, as current GC declines with increased chromosome length (Spearman’s *ρ* = -0.4483, p < 0.0001), but the equilibrium GC* is not related to chromosome length (Spearman’s *ρ* = 0.1436, p = 0.1457). These results indicate that smaller chromosomes have been subject to stronger GC accumulation in the past, perhaps due to higher telomeric recombination rate on small chromosomes causing stronger GC-biased gene conversion. The telomere of the human chromosome 2 (red point in Figure 4C) is an outlier to this trend, as this chromosome is a human-specific fusion and its telomeres likely still retain a stronger deviation from composition equilibrium characteristic to their pre-fusion status as parts of the shorter 2A and 2B chromosomes (still present in other great apes). We conducted a linear regression analysis with GC difference as the response variable and the other three parameters as explanatory variables (Table 2). Every explanatory variable had a significant effect, and together they explain ∼68% of the variance in the deviation of telomeric nucleotide composition from equilibrium.

**Figure 4.**
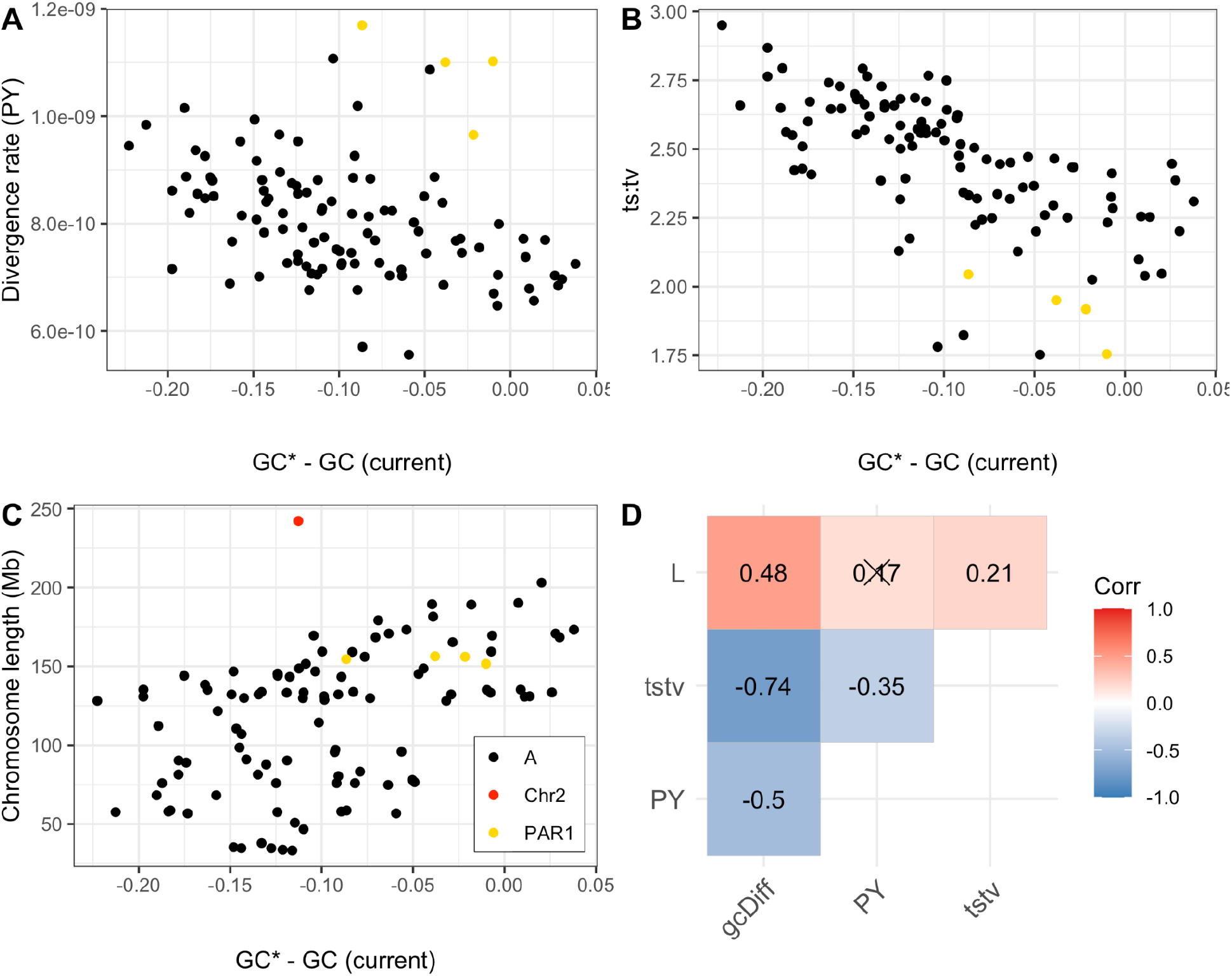
Correlation of the difference between equilibrium GC proportion (GC*) and current GC proportion with **A**. per site and year (PY) divergence rates, **B**. the ts:tv ratios and **C**. chromosome lengths (a telomere of the human chromosome 2 is highlighted in red). **D**. Spearman’s partial correlation coefficients for all pairwise combinations of the three parameters: difference between equilibrium GC* proportion and current GC (gcDiff), per site and year divergence rate (PY), the ts:tv ratio (tstv) and chromosome length (L). Crossed-out coefficients are non-significant (*p* > 0.05). For each telomere we consider four values corresponding to the four external branches of the great ape phylogeny. In total, we consider N=100 estimates for autosomes (across 25 alignable telomeres) and N=4 estimates for PAR1.

**Table 2.** Linear regression model with the response variable as the deviation from nucleotide composition equilibrium (GC*-GC) and explanatory variables as the per site and year divergence rate, the transition:transversion ratio and chromosome length (GC*-GC ∼ div. rate + ts:tv + length). For each telomere we consider four values corresponding to the four external branches of the great ape phylogeny. In total, we consider N=100 estimates for autosomes (across 25 alignable telomeres) and N=4 estimates for PAR1.

|  | Estimate | Standard error | p-value |
| --- | --- | --- | --- |
| (Intercept) | 0.4309 | 0.0506 | $1.7317 \times 10^{-13}$ |
| div. rate | -223.2882 | 30.6623 | $7.6641 \times 10^{-11}$ |
| ts:tv | -0.1611 | 0.0145 | $4.1528 \times 10^{-19}$ |
| length (Mb) | 0.0004 | $8.0120 \times 10^{-5}$ | $1.9489 \times 10^{-6}$ |
| $R^2 = 0.6774$ ($p < 2.2 \times 10^{-16}$) | | | |

### Substitution spectra of telomeres are similar across great apes

We next divided substitutions into seven classes, including a CpG→TpG class, and calculated the count proportion of each class (*i*.*e*., substitution spectrum) for every telomere and great ape species (Figure 5A). We calculated the difference between substitution proportions and the corresponding mean proportion (specific to a species and substitution class), normalized by the standard deviation (Z-score), for all telomeres across all species and substitution classes (Figure 5B). Interestingly, PAR1 and telomere 8L have a similar pattern of proportion differences characterized most strongly by an excess of C→G transversions. This similarity between the two telomeres likely stems from two distinct sources, as the C→G mutagenic signature has been associated with male meiotic double-strand breaks on the X chromosome in humans [22], while telomere 8L has been shown to be a female *de novo* mutation (DNM) hotspot, with a maternal C→G mutation rate that is 50-fold greater than the genome average [66]. Furthermore, non-human great ape species also exhibit the C→G substitution excess on 8L and PAR1, indicating that these mutation hotspots are conserved across the great ape phylogeny. In addition to an excess of C→G transversions, 8L and PAR1 have a general decrease in transition and increase in transversion proportions, resulting in the lowest ts:tv ratios among all telomeres (Supplementary table 1). Telomeres 16L and 16R have also been identified as DNM hotspots and, while they do show an excess of C→G transversions, their ts:tv ratios are similar to those of other autosomal telomeres. The higher similarity between 8L and PAR1 may stem from the dependence of telomere substitution dynamics on divergence rates, the ts:tv ratio and chromosome length (Figure 4; Table 2) - indeed, chromosome 8 and X are very similar with respect to all three parameters (Supplementary table 1).

**Figure 5.**
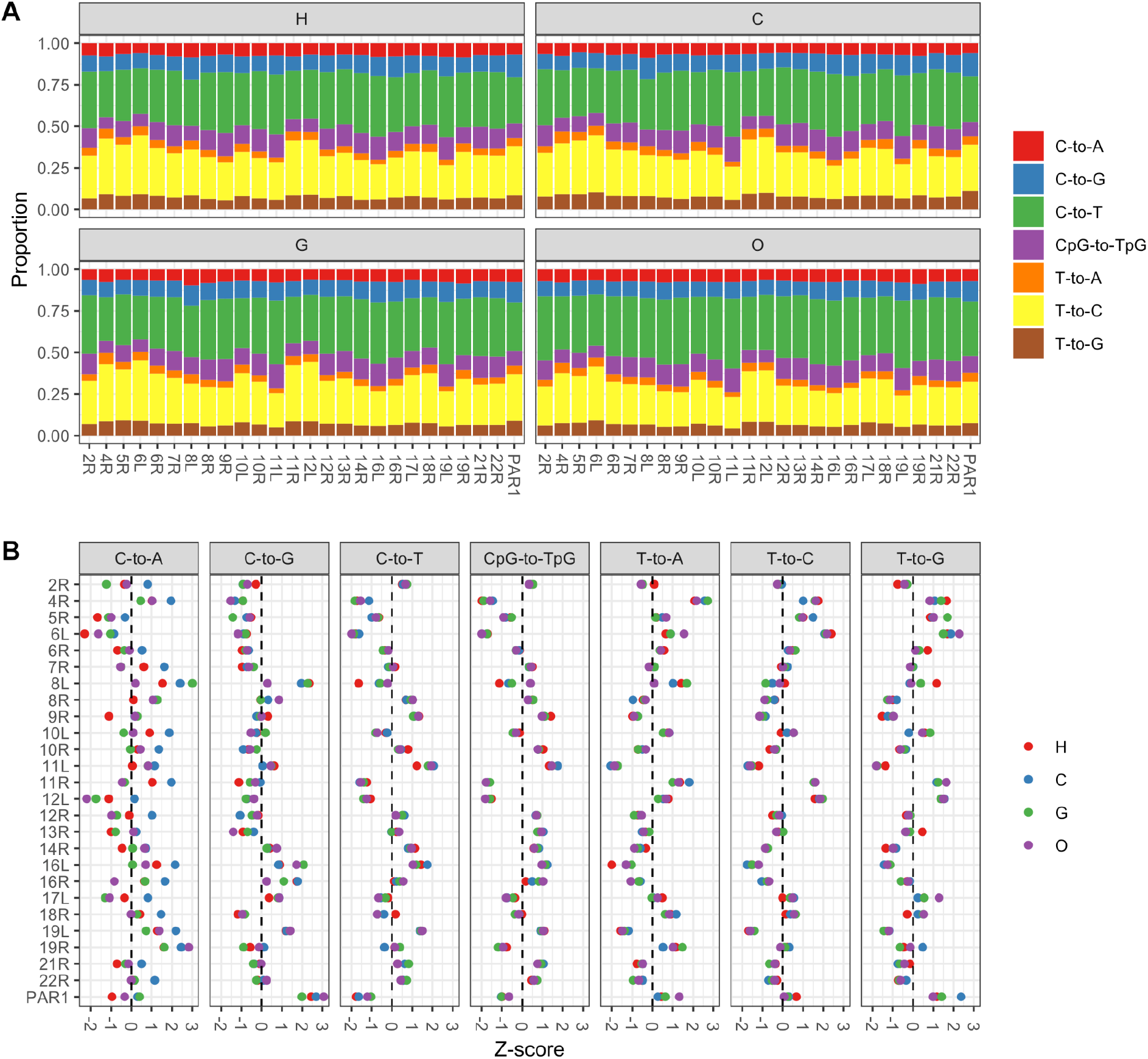
**A**. Substitution spectra for human (H), chimpanzee (C), gorilla (G) and orangutan (O) across 25 autosomal telomeres and PAR1. **B**. Difference between relative substitution proportions and mean relative proportions normalized by the standard deviation (Z-score) for each substitution class, telomere and species. For each telomere and substitution class, we plot four values corresponding to the four external branches of the great ape phylogeny. In total, we plot N=700 estimates for autosomes (across 25 alignable telomeres) and N=28 estimates for PAR1.

### Recombination and diversity in PAR1

In the absence of family data, we used the LDhat program to generate PAR1 recombination rates (*ρ* = 4*N*_*e*_*r*; where *r* is the rate in Morgans) for eight subspecies of great apes, to compare with the published human PAR1 map, inferred from directly observed crossovers in human pedigrees [21] (Figure 6A). The LDhat recombination rate estimates represent population-wide sex-averages, and were converted into cM/Mb by assuming a sex-averaged recombination rate of 9.01 cM/Mb, as inferred in the human PAR1 [21]. Notably, across the subspecies studied, the recombination rate per physical distance is consistently high towards the telomeric end of PAR1, though humans have a significant uptick close to the pseudoautosomal boundary. The majority of recombination events occur between positions 0.25-1.25 Mb of PAR1 with the strongest peak of recombination inferred for *P. troglodytes ellioti*, located between base pairs 330,001-340,000. The peak has an extremely high recombination rate of 90.42 cM/Mb and overlaps the gene *PPP2R3B*, known to be a recombination hotspot [67], and highly expressed during spermatogenesis [68]. We next lifted over genomic coordinates of non-human species to the human reference and estimated the between-species correlation of recombination rates for 10 kb genomic windows (Figure 6B). Generally, correlation coefficients between maps of the different species are positive. The correlations are strongest between species of the same genus; among subspecies of the *Pan* genus, the overall strongest inferred correlation is between the *P. troglodytes ellioti* and *P. troglodytes schweinfurthii* maps (Spearman’s *ρ* = 0.6739, *p* < 0.0001), and the *P. abelli* and *P. pygmaeus* map correlation is stronger than the correlations with non-orangutan species (Spearman’s *ρ* = 0.5752, *p* < 0.0001). Taken together, these results imply that, while recombination in PAR1 is broadly similar across great apes, genus-specific patterns of PAR1 recombination are also observable at the studied phylogenetic scale.

**Figure 6.**
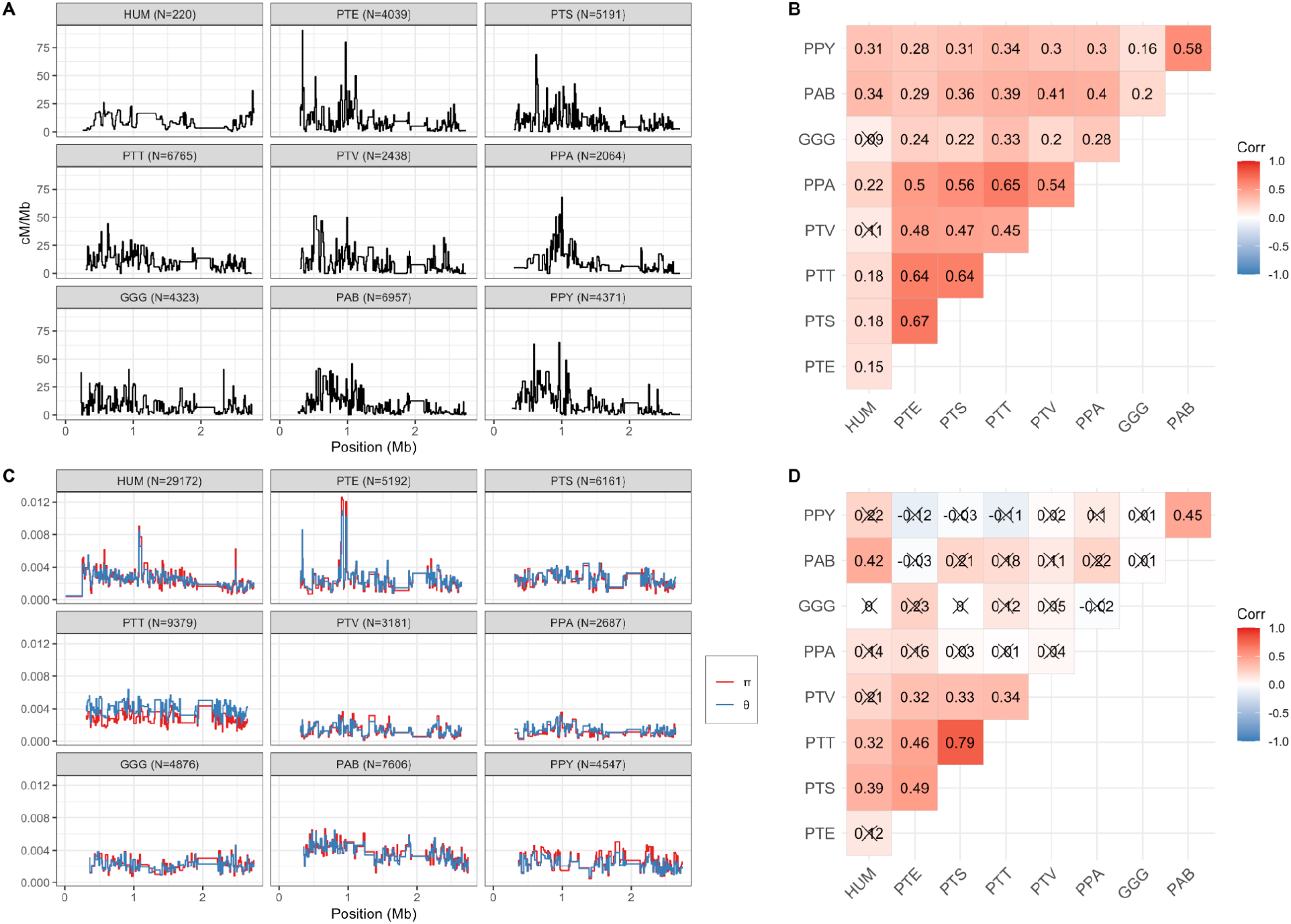
**A**. Sex-averaged recombination rates for 10 kb windows of PAR1 in humans (HUM) and eight subspecies of great apes (PTE = *P. troglodytes ellioti*; PTS = *P. troglodytes schweinfurthii*; PTT = *P. troglodytes troglodytes*; PTV = *P. troglodytes verus*; PPA = *P. paniscus*; GGG = *G. gorilla gorilla*; PAB = *P. abelii*; PPY = *P. pygmaeus*). Numbers of polymorphic sites used to infer recombination maps are presented in the parentheses. **B**. Spearman’s coefficients for correlations between recombination maps at the 10 kb scale. **C**. Nucleotide diversity for 10 kb windows of PAR1 in humans and great apes, measured as *π* and Watterson’s *θ*. Numbers of polymorphic sites used to infer diversity statistics are presented in the parentheses. **D**. Spearman’s coefficients for correlations between nucleotide diversity *π* at the 10 kb scale. Crossed-out coefficients are non-significant (*p* > 0.05). We only consider 10 kb regions with more than 2,500 callable sites.

Nucleotide diversity in PAR1 is on average 2.3 times higher than autosomal estimates (Table 3), and follows the recombination pattern, with elevated rates close to the telomere end (Figure 6C), and a strong within-genus correlation (Figure 6D). The correlation between recombination and average pairwise diversity *π* is positive for all species (Figure 7), consistent with recombination-associated mutagenesis [22,30,31].

**Table 3.** Estimates of PAR1 population genetic parameters for human (HUM) and great ape populations (PTE = *P. troglodytes ellioti*; PTS = *P. troglodytes schweinfurthii*; PTT = *P. troglodytes troglodytes*; PTV = *P. troglodytes verus*; PPA = *P. paniscus*; GGG = *G. gorilla gorilla*; PAB = *P. abelii*; PPY = *P. pygmaeus*).

| | $\rho$ | $\pi$ | $\theta_{\text{PAR}} (\theta_A)$ | Tajima's D | $N_e$ | $\mu$ (PG) | $\mu$ (PY) | B |
| --- | --- | --- | --- | --- | --- | --- | --- | --- |
| HUM | - | 0.0025 | 0.0025 (0.0009) | -0.0541 | - | - | - | 0.6745 |
| PTE | 12,922 | 0.0024 | 0.0026 (0.0013) | -0.2500 | 14,138 | $4.31 \times 10^{-8}$ | $1.72 \times 10^{-9}$ | 0.6801 |
| PTS | 16,141 | 0.0023 | 0.0026 (0.0016) | -0.3273 | 17,661 | $3.31 \times 10^{-8}$ | $1.32 \times 10^{-9}$ | 0.6064 |
| PTT | 32,855 | 0.0028 | 0.0039 (0.0023) | -1.0783 | 35,948 | $1.97 \times 10^{-8}$ | $0.79 \times 10^{-9}$ | 0.8125 |
| PTV | 9,136 | 0.0013 | 0.0014 (0.0008) | -0.2132 | 9,996 | $3.33 \times 10^{-8}$ | $1.33 \times 10^{-9}$ | 0.7302 |
| PPA | 15,293 | 0.0011 | 0.0014 (0.0005) | -0.7477 | 16,732 | $1.70 \times 10^{-8}$ | $0.68 \times 10^{-9}$ | 0.7704 |
| GGG | 12,584 | 0.0023 | 0.0022 (0.0016) | 0.1314 | 13,769 | $4.10 \times 10^{-8}$ | $2.16 \times 10^{-9}$ | 0.7640 |
| PAB | 11,901 | 0.0038 | 0.0037 (0.0019) | 0.1374 | 13,021 | $7.26 \times 10^{-8}$ | $2.79 \times 10^{-9}$ | 0.6531 |
| PPY | 13,728 | 0.0027 | 0.0023 (0.0013) | 0.6627 | 15,020 | $4.48 \times 10^{-8}$ | $1.72 \times 10^{-9}$ | 0.4962 |

$\rho = 4N_e r$ of the whole PAR1 region, as measured by LDhat
$\pi = 4N_e \mu$ ; average pairwise diversity estimator of PAR1
$\theta = 4N_e \mu$ ; Watterson's diversity estimator ( $\theta_{\text{PAR}} = \text{PAR1 } \theta$ ; $\theta_A = \text{autosomal } \theta$ )
$N_e$ = effective size of the population
$\mu$ = per site and per generation (PG), or per year (PY) mutation rate
$B = 4N_e b$ ; strength of GC-biased gene conversion in PAR1

**Figure 7.**
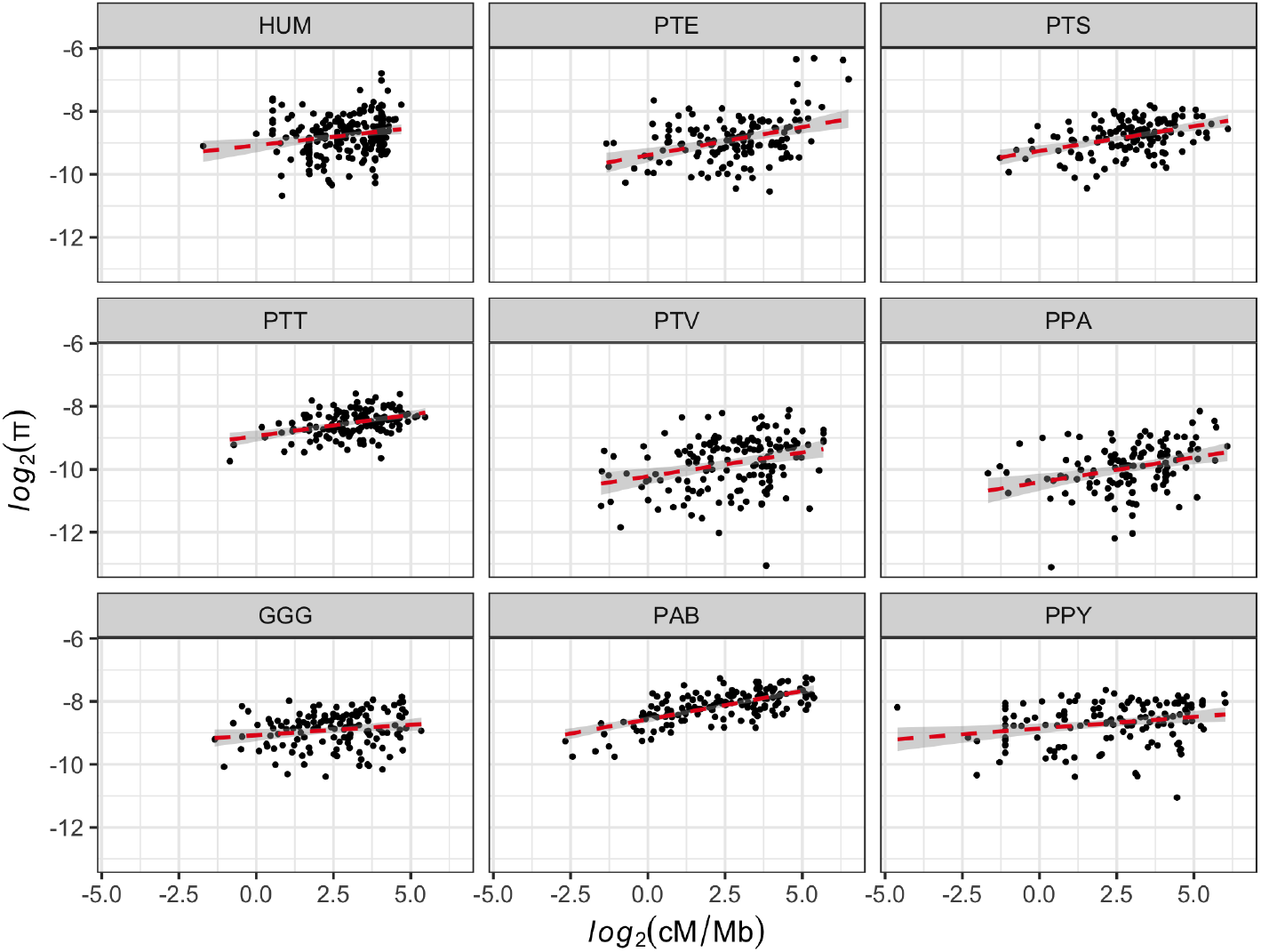
Correlation of recombination rate and nucleotide diversity, measured as *π*, for 10 kb windows of PAR1 in humans (HUM) and eight subspecies of great apes (PTE = *P. troglodytes ellioti*; PTS = *P. troglodytes schweinfurthii*; PTT = *P. troglodytes troglodytes*; PTV = *P. troglodytes verus*; PPA = *P. paniscus*; GGG = *G. gorilla gorilla*; PAB = *P. abelii*; PPY = *P. pygmaeus*). We only consider 10 kb regions with more than 2,500 callable sites.

Given LDhat estimates of *ρ* (4*N*_*e*_*r*) for PAR1, where *r* = cM/100, and assuming that the sex-averaged genetic length of PAR1 in great apes is equal to the human estimate of 22.85 cM [21], we can calculate the effective size (*N*_*e*_) for each population (Table 3). For five out of eight great ape species, *N*_*e*_ estimates fall within the previously inferred ranges [69]. For *P. paniscus, N*_*e*_ is higher than the previously inferred upper limit (16,732 *vs*. 10,800 individuals), while for *G. gorilla gorilla* and *P. abelii, N*_*e*_ is lower than the previously inferred lower limit (13,769 *vs*. 20,132 and 13,021 *vs*. 16,731 individuals, respectively). We can similarly calculate the PAR1-specific per generation mutation rate *μ* by taking the ratio *π*/*ρ* = *μ*/r (Table 3). We also report *μ* as a per year rate by assuming generation times of 25 years for chimpanzees, 19 years for the gorilla and 26 years for the orangutan. Compared to the per year mutation rate estimated from great ape trios (0.64 ×10^−9^; [61]), PAR1 yearly *μ* is on average 2.44-fold larger, ranging from a 1.06-fold increase in *P. paniscus* to a 4.36-fold increase in *P. abelii*. While this mutation range overlaps the per year divergence range of PAR1 (Table 1), mutations notably occur at a higher yearly rate compared to substitutions, indicating that extant nucleotide diversity levels do not fully reflect the long-term substitution dynamics of PAR1.

### Differences between segregating polymorphisms and substitution spectra in PAR1

We next explore the differences between PAR1 diversity and substitution spectra, and their dependence on the recombination rate. We therefore divided the PAR1 sequence of each great ape population into 10 kb windows with low (≤ 4.25 cM/Mb), medium (4.25 < cM/Mb ≤ 11.29) and high recombination rates (> 11.29 cM/Mb), based on recombination rate terciles determined across all maps of the nine populations studied (Figure 6A). Since the substitution spectrum estimates are based on the reference alignment, *i*.*e*., all sequences within the alignment are considered simultaneously, while the division of PAR1 regions into recombination bins is population-based, we applied the following protocol when inferring the substitution spectrum of a specific genus and recombination bin. Due to the strong within-genus correlation of recombination rates (Figure 6B), we inferred recombination maps for each genus by calculating the mean recombination rate for each 10 kb region across all its sampled populations. For the human and gorilla datasets, with one population per genus, we simply used the inferred rates. Based on each genus-specific map, we categorised 10 kb regions into the three recombination bins to infer divergence rates and posterior substitution counts using the phyloFit program. In total, this procedure results in four different divisions of PAR1 (one for each genus), each with three genus-specific recombination bins (*i*.*e*., 12 distinct combinations of PAR1 10 kb regions in total). Table 4 reports divergence rates for the different recombination bins. Notably, divergence rates increase with increasing recombination rates for all lineages studied. The increase in divergence between the low and high recombination bins for extant species ranges from 1.24-fold in gorillas to 1.54-fold in orangutans. In Figure 8 we show the substitution and extant diversity spectra for PAR1 across the three recombination bins. The most notable difference between the two spectra is the decrease of CpG→TpG substitutions compared to extant segregating variation. This trend is observable across all three recombination bins. Furthermore, for all substitution and diversity spectra, the C→G proportion is higher in the highest, compared to the lowest recombination bin, in line with the observation that this mutation type is associated with male meiotic double-strand breaks on the X [22].

**Table 4.**
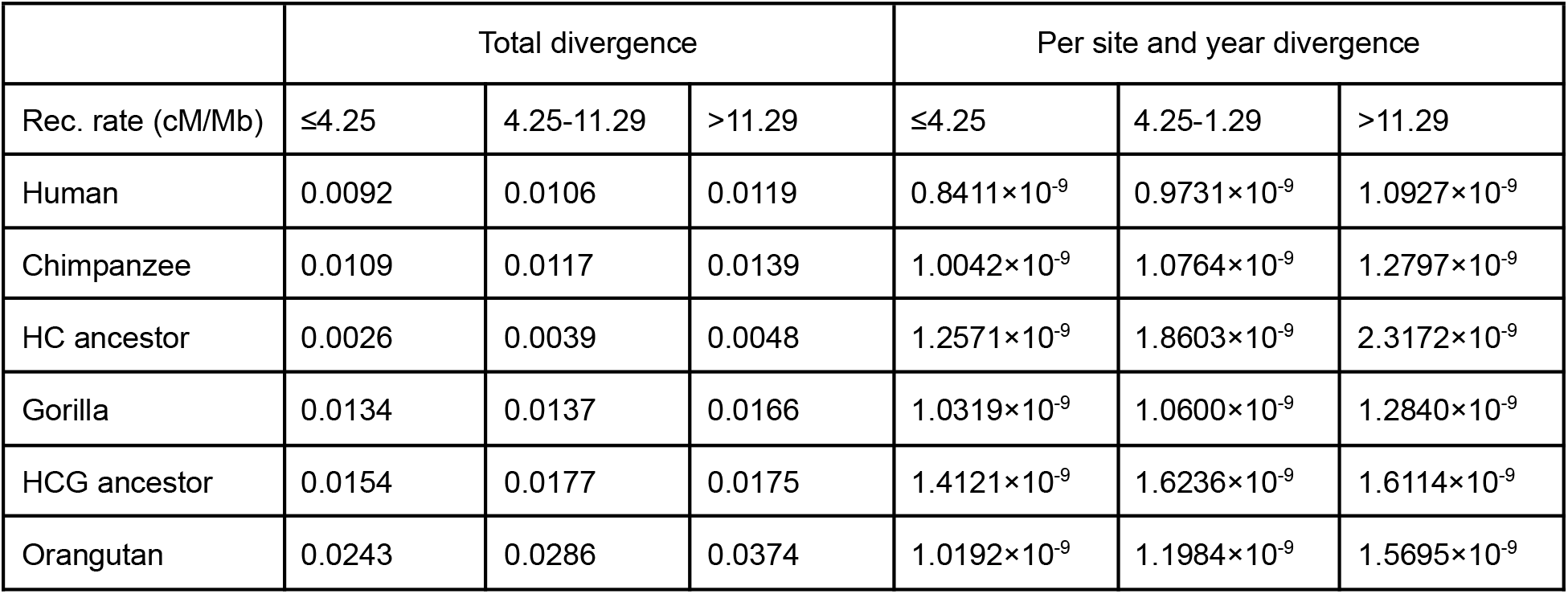
Divergence estimates based on phyloFit estimation and divergence times from [61] for the PAR1 sequence, across three recombination rate bins.

|  | Total divergence |  |  | Per site and year divergence |  |  |
| --- | --- | --- | --- | --- | --- | --- |
| Rec. rate (cM/Mb) | ≤4.25 | 4.25-11.29 | >11.29 | ≤4.25 | 4.25-11.29 | >11.29 |
| Human | 0.0092 | 0.0106 | 0.0119 | $0.8411 \times 10^{-9}$ | $0.9731 \times 10^{-9}$ | $1.0927 \times 10^{-9}$ |
| Chimpanzee | 0.0109 | 0.0117 | 0.0139 | $1.0042 \times 10^{-9}$ | $1.0764 \times 10^{-9}$ | $1.2797 \times 10^{-9}$ |
| HC ancestor | 0.0026 | 0.0039 | 0.0048 | $1.2571 \times 10^{-9}$ | $1.8603 \times 10^{-9}$ | $2.3172 \times 10^{-9}$ |
| Gorilla | 0.0134 | 0.0137 | 0.0166 | $1.0319 \times 10^{-9}$ | $1.0600 \times 10^{-9}$ | $1.2840 \times 10^{-9}$ |
| HCG ancestor | 0.0154 | 0.0177 | 0.0175 | $1.4121 \times 10^{-9}$ | $1.6236 \times 10^{-9}$ | $1.6114 \times 10^{-9}$ |
| Orangutan | 0.0243 | 0.0286 | 0.0374 | $1.0192 \times 10^{-9}$ | $1.1984 \times 10^{-9}$ | $1.5695 \times 10^{-9}$ |

**Figure 8.**
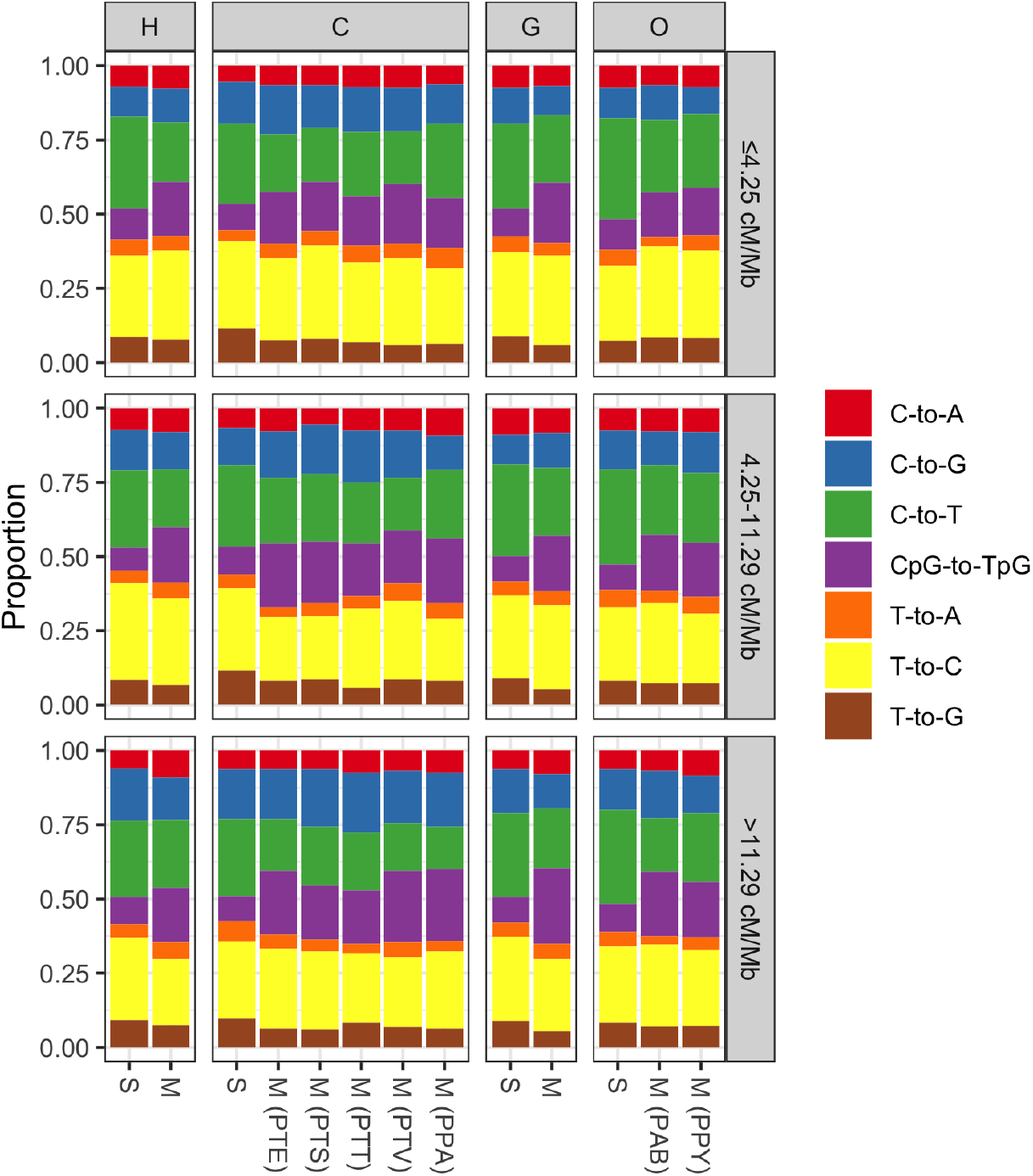
Substitution and diversity spectra for the human (H), chimpanzee (C), gorilla (G) and orangutan (O) PAR1, across three recombination bins (rows). Substitution and diversity spectra are denoted by S and M, respectively. Multiple diversity spectra are plotted for chimpanzees and orangutans, each corresponding to a particular subspecies (PTE = *P. troglodytes ellioti*; PTS = *P. troglodytes schweinfurthii*; PTT = *P. troglodytes troglodytes*; PTV = *P. troglodytes verus*; PPA = *P. paniscus*; PAB = *P. abelii*; PPY = *P. pygmaeus*), while the diversity spectra for humans and gorillas correspond to the human YRI and *G. gorilla gorilla* populations, respectively.

Figure 8 indicates that CpG mutations are the main contributors to the discrepancy between long-term substitution rates and segregating polymorphism. Such an increase in extant diversity compared to the long-term neutral expectation (*i*.*e*., substitution spectrum) is expected when a fixation bias favouring GC nucleotides is opposed by AT-biased mutation [70]. In humans and great apes, the preferred fixation of GC nucleotides is mediated by recombination, and is termed GC-biased gene conversion (gBGC; [33–35]). To assess the strength of gBGC in PAR1, we constructed allele frequency spectra (AFS) based on GC frequency categories of segregating variants and inferred the bias parameter *B* = 4*N*_*e*_*b*, where *b* is a bias coefficient [36]. For this analysis, we omitted sites segregating for CpG transversions, due to their idiosyncratic gBGC dynamics [71], and calculated *B* only for the other sites (Table 3), and separately for three mutation categories: non-CpG and CpG transitions, and non-CpG transversions (Figure 9). Generally, PAR1 *B* estimates are approximately twice the autosomal estimates [71,72], reflecting the high PAR1 recombination rate, with CpG sites experiencing the strongest gBGC dynamics. Therefore, the exceptionally strong gBGC at CpG sites likely contributes to the paucity of this mutation class in the substitution spectrum as CpG→TpG mutations are ultimately fixed back as GC nucleotides, while extant diversity at these sites is elevated due to the opposition between exceptionally strong GC→AT mutation and AT→GC fixation forces [70].

**Figure 9.**
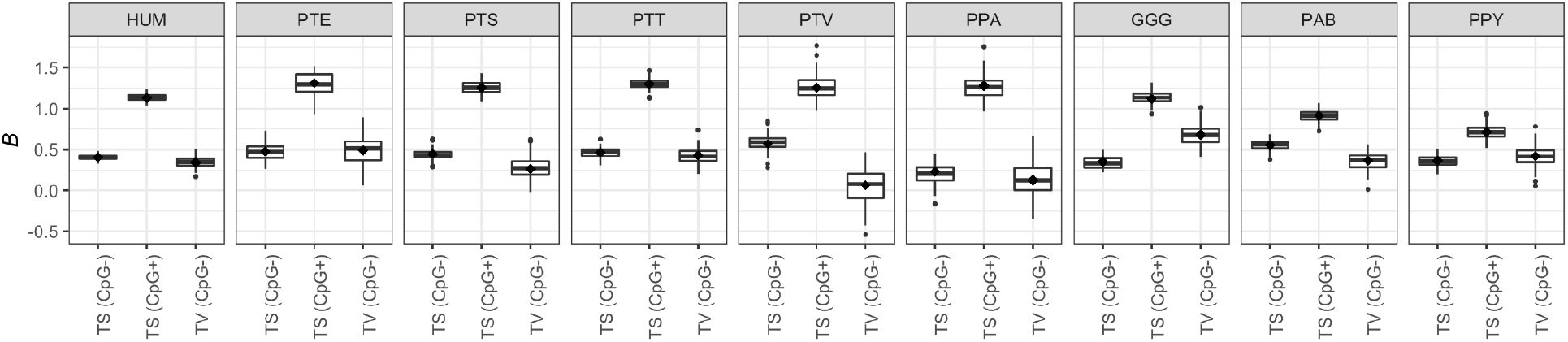
PAR1 *B* estimates for non-CpG transitions (TS; CpG–), CpG transitions (TS; CpG+), and GC-changing transversions (TV; CpG–) for the human YRI population (HUM) and eight subspecies of great apes (PTE = *P. troglodytes ellioti*; PTS = *P. troglodytes schweinfurthii*; PTT = *P. troglodytes troglodytes*; PTV = *P. troglodytes verus*; PPA = *P. paniscus*; GGG = *G. gorilla gorilla*; PAB = *P. abelii*; PPY = *P. pygmaeus*). Distributions of *B* estimates were obtained from 100 bootstrapped allele frequency spectra for each mutation type and subspecies. The “diamond” points correspond to the maximum likelihood *B* estimate.

### No evidence for sexually antagonistic alleles in PAR1

As regions closely linked to the pseudoautosomal boundary can potentially maintain polymorphisms for sexually antagonistic (SA) alleles under a wider parameter range than other genome regions, we looked for differences in sex-specific allele frequencies. In Figure 10A, we show absolute frequency differences between females and males for all segregating sites and species. Frequency differences for the human YRI population reach about ∼25%, and up to 100% for great ape populations. However, when applying Fisher’s exact tests to assess the significance of frequency differences, followed by Bonferroni’s correction for multiple testing, we found no significant frequency differences in any of the populations (Figure 10B). Due to the small sample sizes for non-human populations, the significance threshold is unreachable for most of these populations, as demonstrated by a SNP at position 180,684 for the *P. paniscus* population. Although this SNP is fixed for different alleles in females and males, the frequency difference is non-significant after *p*-value correction. In the human YRI population, the SNP that is closest to a significant frequency difference is located at position 1,322,855 (rs146912994) in the intron of the *CSF2RA* gene. The T allele of this SNP segregates at 15.69% frequency in males, while females are fixed for the C allele. Furthermore, this SNP is not present in Eurasian populations [73]. However, according to the dbSNP database (https://www.ncbi.nlm.nih.gov/), there are currently no phenotypes associated with this SNP that would imply sexually antagonistic evolution.

**Figure 10.**
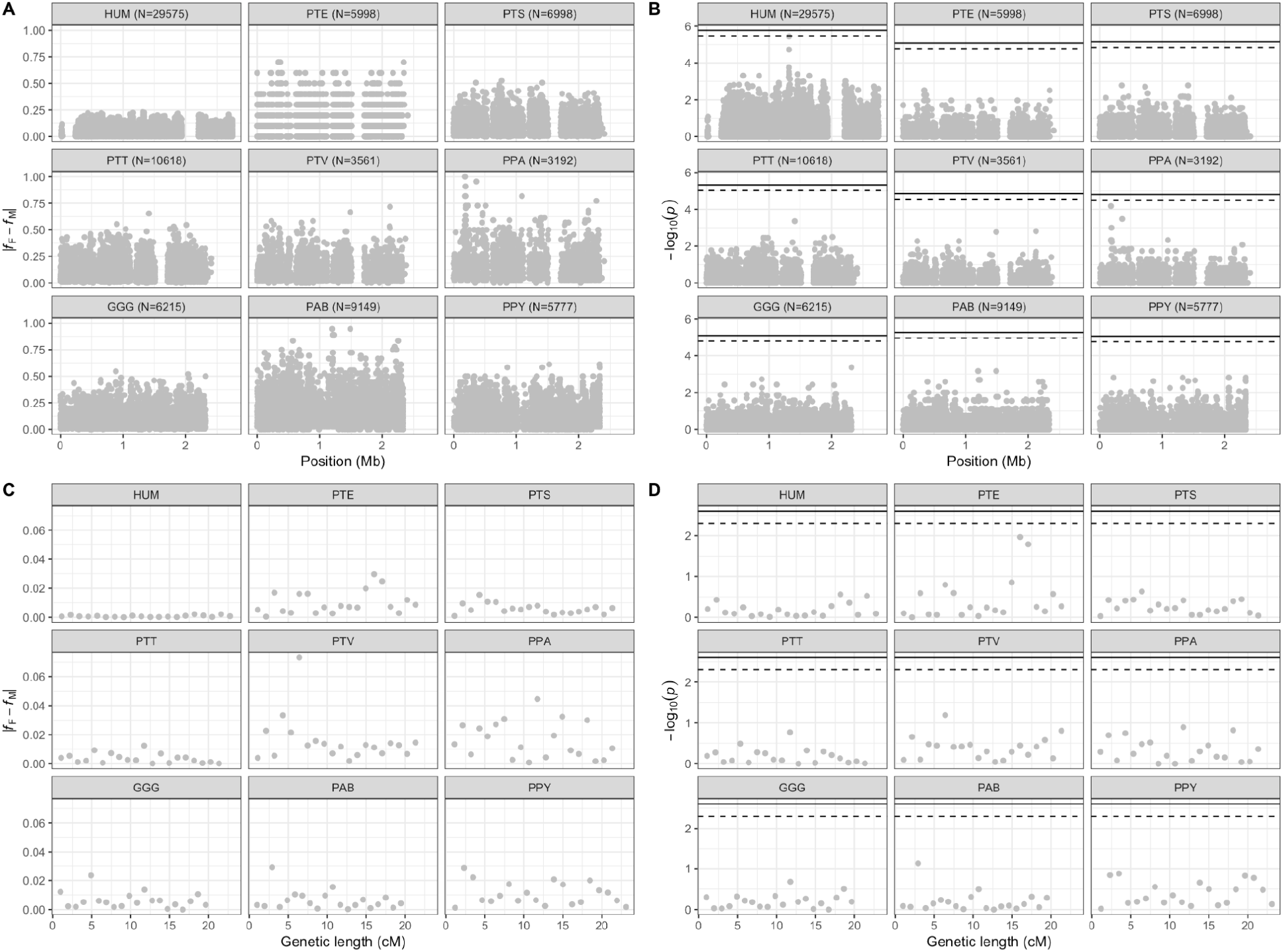
**A**. Absolute differences between female (*f*_*F*_) and male (*f*_*M*_) allele frequencies for sites segregating in humans (HUM) and eight subspecies of great apes (PTE = *P. troglodytes ellioti*; PTS = *P. troglodytes schweinfurthii*; PTT = *P. troglodytes troglodytes*; PTV = *P. troglodytes verus*; PPA = *P. paniscus*; GGG = *G. gorilla gorilla*; PAB = *P. abelii*; PPY = *P. pygmaeus*). Numbers of polymorphic sites plotted in each panel are presented in the parentheses. **B**. *p*-values of Fisher’s exact tests for between-sex allele frequency differences. **C**. Window-based absolute differences between female and male allele frequencies for sites segregating in 20 windows of approximately 1 cM in genetic length in humans and eight subspecies of great apes. **D**. Window-based *p*-values of Fisher’s exact tests for average between-sex allele frequency differences. The solid and dashed horizontal lines in panels **B** and **D** represent the significance thresholds at *p* < 0.05 and *p* < 0.1, respectively, after Bonferroni’s correction for multiple testing.

Since PAR1 segregates for a large number of sites, we were able to test whether frequency differences increase with proximity to the pseudoautosomal boundary by the following analysis, as would be expected in the presence of SA selection [6]. We separated segregating variants into 20 non-overlapping windows, each approximately 1 cM in genetic length, and calculated window-based average differences between female and male allele frequencies. We find no evidence for such a SA gradient (Figure 10C), nor generally any significant differences between sex-specific allele frequencies when applying Fisher’s exact test to the sums of female and male allele counts in each window (Figure 10D).

### Evolution of PAR1 during recent hominid speciation

We next analyze archaic PAR1 sequences of a Neanderthal and Denisovan individual. In total, 251,571 putatively neutral base pairs were alignable between the archaic sequences and the human and chimpanzee reference sequences. Pairwise divergence between the human and archaic samples are 0.0021, 0.0022 and 0.0019 for human-neanderthal, human-denisovan and neanderthal-denisovan pairs, respectively. Figure 11 shows a neighbor-joining tree of archaic individuals and the human reference, using the chimpanzee sequence as an outgroup. The Neanderthal and Denisovan individuals are grouped into a separate clade, as previously observed for phylogenetic trees based on autosomal data [50,74]. Therefore, while a recent study showed that the Y chromosome introgressed from humans into neanderthals [49], our results indicate that the neanderthal PAR1 retained the autosomal-like phylogenetic relationship (*i*.*e*., avoided partial replacement with the invading human Y-linked PAR1).

**Figure 11.**
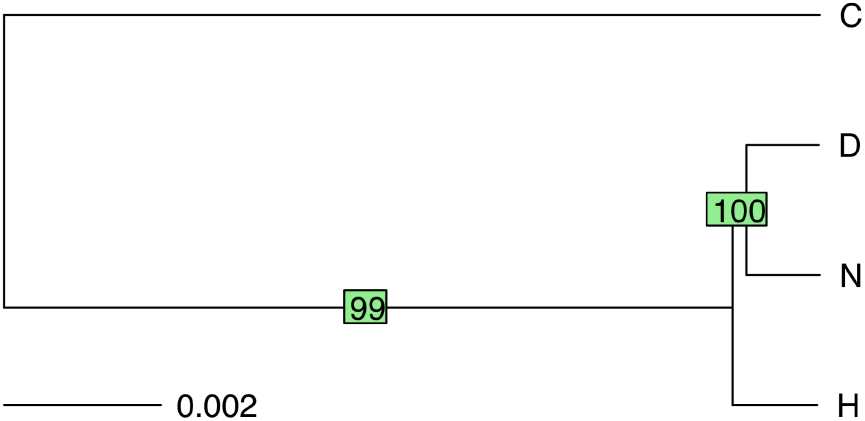
PAR1 neighbor-joining tree for human (H), Neanderthal (N), Denisovan (D) and chimpanzee (C) sequences. Internal branch supports of 100 bootstrap samples are shown in green rectangles.

We estimate average nucleotide heterozygosity to be 7.34×10^−4^ and 7.84×10^−4^ per base pair for the Neanderthal and Denisovan sequences, respectively, approximately ∼4× higher than the genome-wide average [50,51]. Figure 12A shows diversity values in 10 kb regions along PAR1. Interestingly, the Denisovan sequence shows high telomeric diversity, as in humans (Figure 6C), while neanderthal diversity is relatively low at the telomeric end. The correlation between diversity patterns is positive between human and Denisovan sequences, and non-significant for the other two comparisons (Figure 12B).

**Figure 12.**
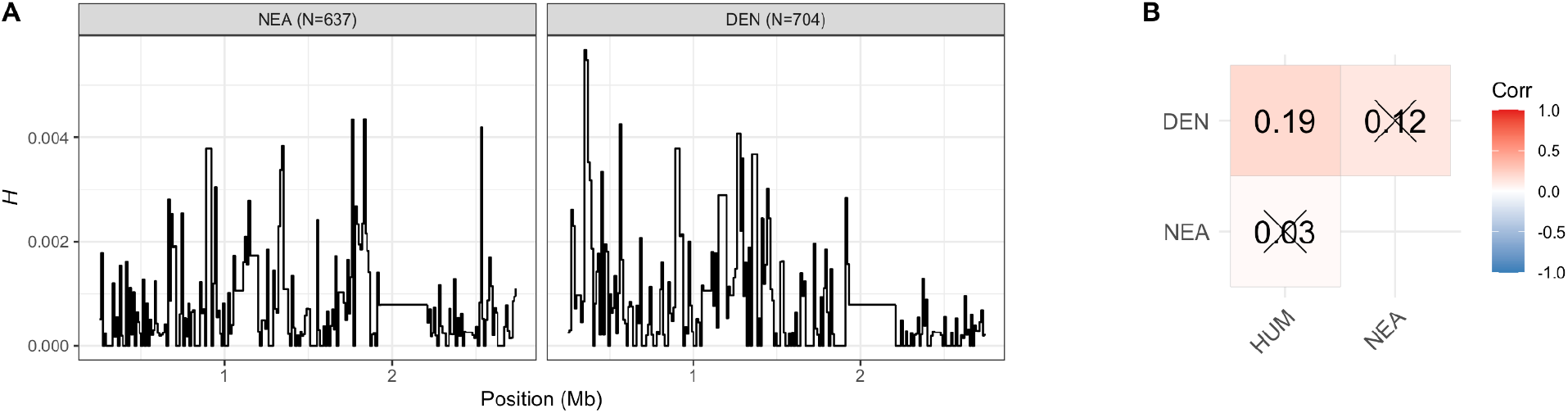
**A**. Nucleotide heterozygosity for 10 kb windows of PAR1 in Neanderthal (NEA) and Denisovan (DEN) sequences, measured as per site heterozygosity *H*. Numbers of polymorphic sites used to infer *H* values are presented in the parentheses. **B**. Spearman coefficients for correlations between diversity at the 10 kb scale, for archaic individuals and humans (HUM). Crossed-out coefficients are non-significant (*p* > 0.05). We only consider 10 kb regions with more than 2,500 callable sites.

The difference in diversity patterns between the archaic sequences could be due to changes in mutational processes during recent *Homo* evolution [48]. Correspondingly, the Denisovan substitution spectrum resembles that of humans more closely than that of Neanderthals (Figure 13). We also generally recover the pattern of relatively higher proportions of C→G transversions and a lack of CpG→TpG transitions in archaics compared to modern humans, as reported in [48]. Additionally, there is a notable excess of C→A transversions in the Neanderthal spectrum, but not in the Denisovan one. This mutation type is enriched in archaic sequences, but non-significantly [48].

**Figure 13.**
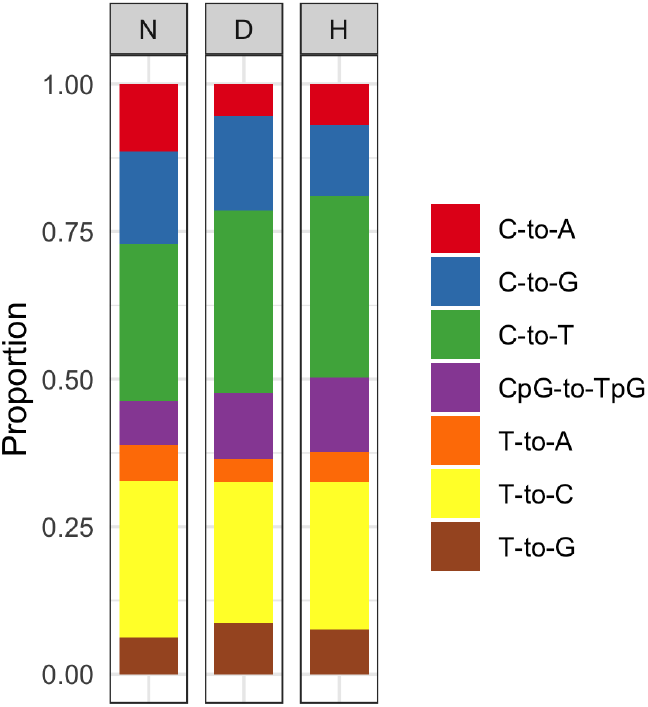
Substitution spectra for the Neanderthal (N), Denisovan (D) and human (H) PAR1.

### Assurance of recombination in PAR1

A recombination event in the pseudoautosomal region during male gametogenesis is crucial for proper sex chromosome segregation and male fertility. In mice, ensuring PAR recombination has been shown to be associated with a unique chromatin structure of the PAR, mediated by repetitive DNA [54]. We therefore sought to characterize repeat elements of the human PAR1, using autosomal telomeres as a reference point. Specifically, we used the Tandem Repeats Finder program [75] to analyse PAR1 and the terminal 3 Mb regions of autosomes, to obtain basic statistics about their repeat content (Figure 14). The largest number of repeat elements was found in PAR1. Furthermore, PAR1 elements usually contained more copies of the consensus pattern, compared with the autosomal regions. In total, almost 25% of PAR1 is composed of repeat sequences - approximately double the proportion of the most repeat-rich autosomal telomeres. Additionally, PAR1 repeats contain the highest proportions of mismatches and indels between adjacent copies. Together, these results imply an unusual repeat structure in the human PAR1, which may aid in recombination assurance, as observed in mice.

**Figure 14.**
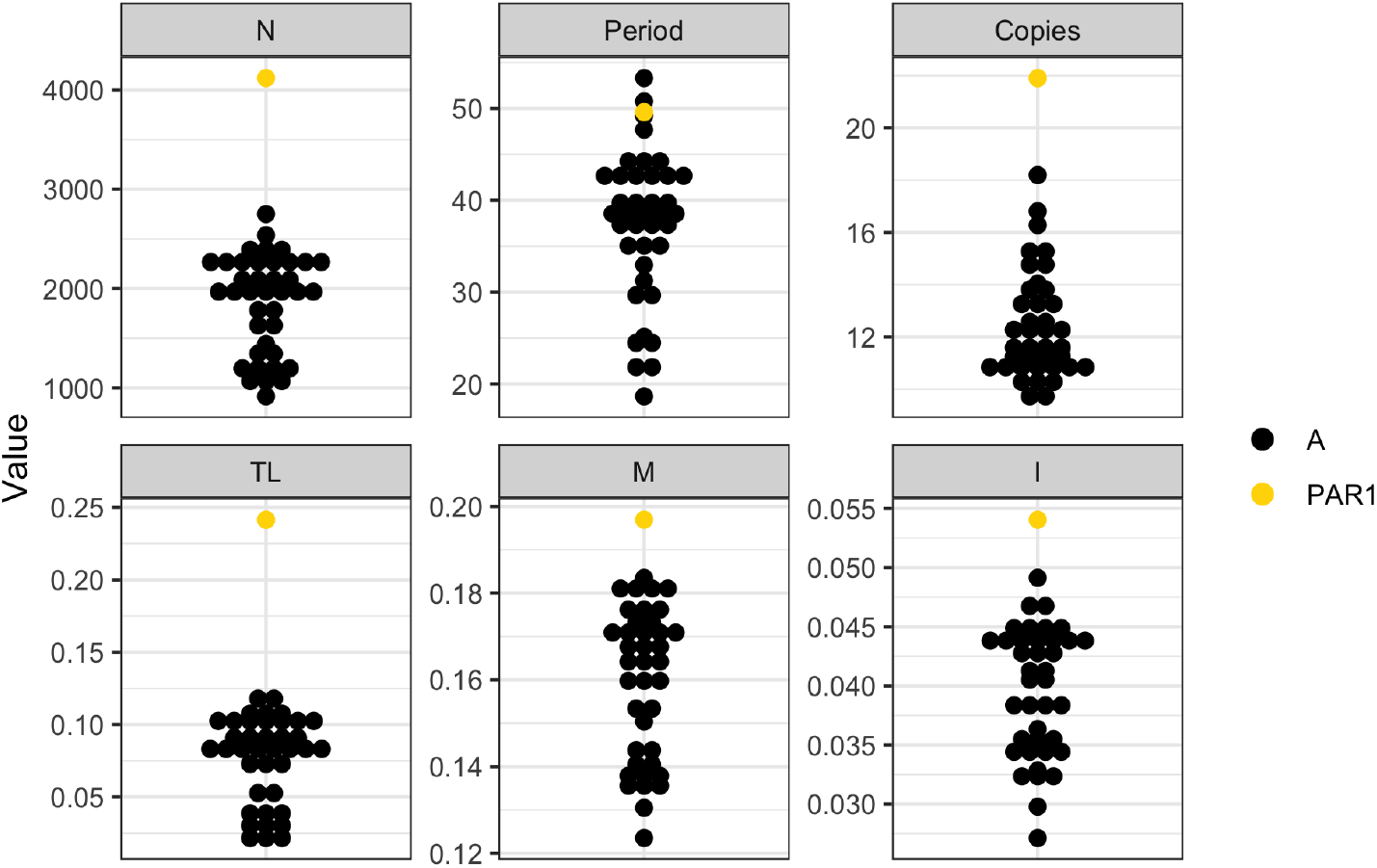
Repeat element statistics for human autosomal (A) telomeres and PAR1. The plotted statistics are as follows: N = the total number of detected repeat elements, Period = the mean period size of the detected repeat elements, Copies = the mean number of copies of the repeating pattern in detected repeat elements, TL = proportion of the total length of the telomere that is covered by repeat elements, M = mean proportion of mismatches between copies of the detected repeat elements, I = mean proportion of indels between copies of the detected repeat elements. In total, we plot 40 points in each panel corresponding to PAR1 and 39 autosomal telomeres (the heterochromatic p-arm telomeres of chromosomes 13, 14, 15, 21 and 22 were excluded from the analysis due to their poor sequence assembly).

Another important determinant of recombination is binding of the recombination motif-recognition protein PRDM9 that mediates meiotic double-strand break (DSB) formation, and was shown to operate in the human PAR1 [55]. We explored the sequence evolution of PAR1 regions within DSB hotspots inferred for individual human males [55]. Specifically, we divided sites of the great ape PAR1 alignment into those falling outside (217,856/239,799; 90.85%) and inside (21,943/239,799; 9.15%) of human DSB hotspots, and ran the phyloFit program to determine divergence rates for the two classes of sites (Table 5). We observe higher divergence of sites within DSB hotspots for the human PAR1, compared with non-hotspot regions, as expected due to the higher rate of sequence evolution within these regions [55]. Interestingly, the same pattern also holds for all other great ape species. Given that the DSB regions were characterized in human males, the fold-increase in divergence is greatest for the human sequence (1.32), as expected, and ranges from 1.14 to 1.3 in the other great ape species. Additionally, substitution spectra for DSB regions show an excess of C→G transversions and a paucity of CpG→TpG transitions compared to non-DSB regions (Figure 15), indicating a stronger mutagenic effect of male recombination, as well as stronger gBGC within these regions, across all great ape species.

**Table 5.** Divergence estimates based on phyloFit estimation and divergence times from [61] for PAR1 regions outside and inside double-strand break (DSB) hotspots as determined by [55].

|  | Outside DSB hotspots | Inside DSB hotspots |
| --- | --- | --- |

|  | Total divergence | Per site and year divergence | Total divergence | Per site and year divergence |
| --- | --- | --- | --- | --- |
| Human | 0.0102 | $0.9373 \times 10^{-9}$ | 0.0135 | $1.2453 \times 10^{-9}$ |
| Chimpanzee | 0.0117 | $1.0724 \times 10^{-9}$ | 0.0152 | $1.3938 \times 10^{-9}$ |
| HC ancestor | 0.0037 | $1.7583 \times 10^{-9}$ | 0.0038 | $1.8105 \times 10^{-9}$ |
| Gorilla | 0.0141 | $1.0859 \times 10^{-9}$ | 0.0162 | $1.2523 \times 10^{-9}$ |
| HCG ancestor | 0.0164 | $1.5087 \times 10^{-9}$ | 0.0185 | $1.7001 \times 10^{-9}$ |
| Orangutan | 0.0275 | $1.1512 \times 10^{-9}$ | 0.0326 | $1.3659 \times 10^{-9}$ |

**Figure 15.**
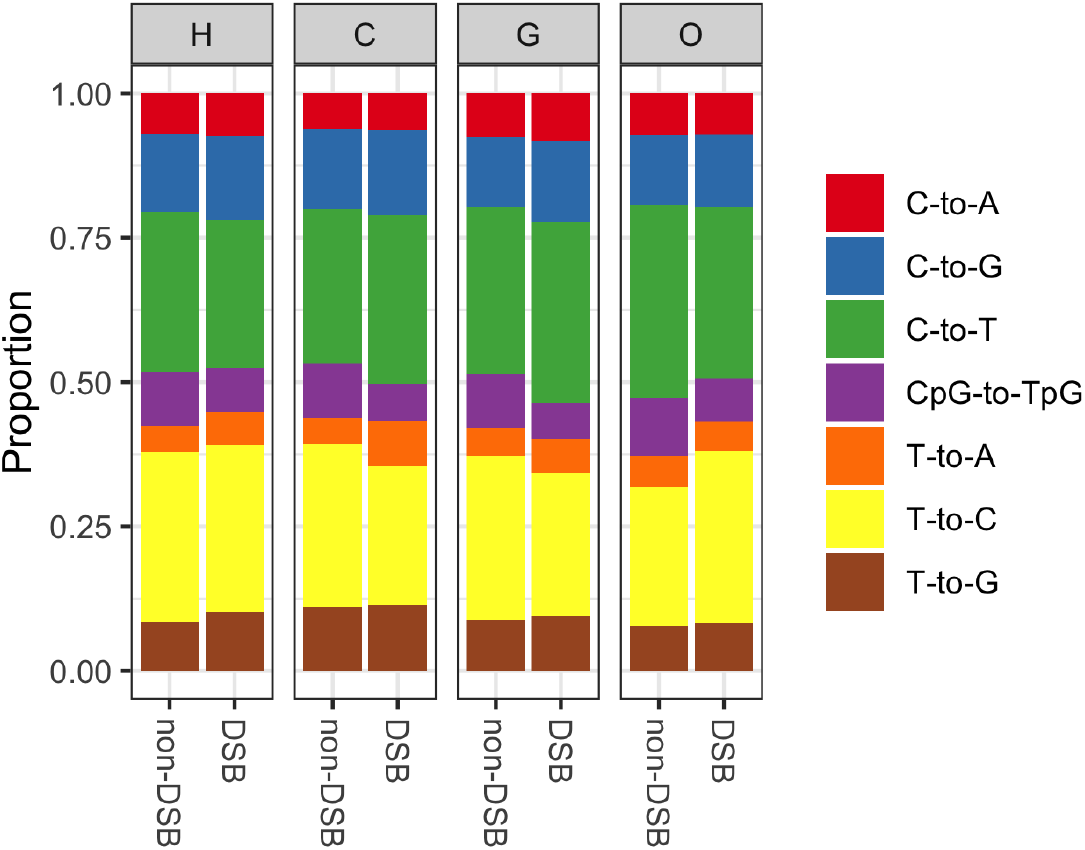
Substitution spectra for the human (H), chimpanzee (C), gorilla (G) and orangutan (O) PAR1 of regions outside and inside of double-strand break hotspots (non-DSB and DSB, respectively).

## Discussion

PAR1 is a unique genomic region due to its linkage to the sex-determining region of the Y chromosome and the status as arguably the most important recombination hotspot in the genome. With this study, we show that PAR1 biology of great apes is mostly governed by the recombination dynamics of this region, rather than antagonistic processes due to sex chromosome linkage.

The nucleotide composition of the human genome is affected by recombination and is known to be evolving towards a lower GC content [34,65]. On the other hand, the GC content of the PAR1 region is relatively close to the equilibrium expectation, compared to similar regions with high male-specific recombination rates (Figure 3). Additionally, PAR1 is closer to nucleotide equilibrium as its predicted equilibrium GC is higher than for autosomal telomeres (Figure 3B). We also show that the deviation from the equilibrium nucleotide content depends on divergence rates, the ts:tv ratio and chromosome length (Figure 4, Table 2). Greater deviations are generally associated with short chromosomes, which may have evolved under relatively stronger gBGC effects, due to higher effective sizes in ancestral populations compared to extant populations [61,65].

As previously observed in humans, we detect a strong enrichment of C→G mutations, associated to male meiotic double-strand breaks on the X [22] in the substitution spectra of all species of great apes (Figure 5). Additionally, the mutation signatures of the telomeric regions of chromosome arms 8L, 16L and 16R [66] are well conserved, indicating a general stability of mutation hotspots and their effects on substitution spectra across the great ape phylogeny. Given the fact that these are not transient hotspots, their ubiquity across a broader set of species would be an interesting avenue of future research.

The analysis of recombination rate and nucleotide diversity revealed expected associations between the two parameters (Figure 6 and 7). Evolution of recombination differences between the genera is evident in PAR1, in line with previous observations of differences between human and chimpanzee recombination maps [76,77]. Additionally, an increased recombination rate close to the pseudoautosomal boundary detected in humans [21] is not seen in other great apes. This may indicate a shift in human PAR1 recombination patterns, but it could also reflect the differences between the methods used to infer the maps (the human map is pedigree-based, whereas the great ape maps are based on LD analyses). An estimation of PAR1 recombination maps using LD-based methods [78,79] for different human populations would be beneficial for a better understanding of this pattern.

The *N*_*e*_ estimates we obtained by using the recombination rate estimate *ρ* (4*N*_*e*_*r*) are fairly accurate for most populations (Table 3). For the *P. paniscus*, PAR1 *N*_*e*_ is larger than previously inferred, while it is lower for the *G. gorilla gorilla* and *P. abelli* populations [69]. This could be due either to processes that cause PAR sequences to evolve differently from autosomal ones, and affect *N*_*e*_ in these populations, or simply due to a limitation of the LDhat method used to infer *ρ*. A more sophisticated method of recombination rate inference using demography-aware models [79] would be valuable in future studies.

Our results quantifying the PAR1-specific mutation and the divergence rates, and their association with recombination (Tables 3 and 4, Figures 6-8), strongly suggest that recombination-associated mutations play a more important role in PAR1 divergence, compared to other genomic regions. In a recent study on the mutagenic impact of recombination, Halldorsson *et al*. (2019) [29] showed that the male *de novo* mutation rate is ∼3.3×10^−8^ per site and generation in the region ±20 kb from male crossovers. If we assume that PAR1 experiences one such crossover per generation and an average background male mutation rate of ∼0.96×10^−8^ per site and generation [29], we estimate that a ∼3.5% increase of PAR1 mutations is due to this one mandatory male crossover event. However, this increase is too small to fully account for the high PAR1-specific *μ* in great apes (Table 3). Therefore, additional mutation inputs, *e*.*g*., from non-crossover events [80] and double strand break (DSB) repair mechanisms [55], are likely to further contribute to a higher mutation rate of the PAR1 sequence.

We also showed that, as expected, the strength of GC-biased gene conversion (Table 3, Figure 9) is higher than for autosomal sequences [71,72,81,82]. By separately estimating the *B* parameters for different classes of mutations, we find that CpG sites are most strongly affected by gBGC dynamics. For gorillas and orangutans, the gBGC strength is somewhat reduced for CpG sites, as observed for autosomes [71]. This mutation class is also underrepresented in the substitution spectrum compared to its frequency in the diversity spectrum (Figure 8). We hypothesize that the discrepancy between the CpG frequency in the diversity and substitution spectrum is due to a somewhat counterintuitive phenomenon [70] that occurs when mutation and fixation forces work in the opposite directions (*i*.*e*., CpG→TpG mutations *vs*. gBGC at CpG-segregating sites). In this case, extant diversity, and therefore the proportion of CpG sites in the diversity spectrum, is elevated beyond the neutral expectation (*i*.*e*., the substitution spectrum). Importantly, this phenomenon may not be limited to CpG sites, and is probably occurring at all GC-changing sites evolving under the extreme PAR1-specific mutation and substitution dynamics. Therefore, the estimated range of PAR1-specific mutation rates *μ* (Table 3) is also likely overestimated, and should be interpreted with caution.

In a recent study, differences between frequencies of PAR1 alleles segregating in females and males have been reported for humans [23]. In contrast to this, our analysis found no alleles that show patterns of sexually antagonistic evolution (Figure 10). We see three possible reasons for this discrepancy. Firstly, for the majority of great ape populations, the sample size is simply too small to reach significance levels, even for sites fixed for different alleles between females and males. Secondly, in the study of human PAR sex-specific allele frequencies, to our knowledge, the authors of [23] did not apply any correction for multiple-testing when assessing frequency differences, which makes their results difficult to reconcile with our own analysis of the human dataset. Thirdly, while we consider only the African YRI population, Monteiro *et al*. (2021) [23] study frequency differences using either all individuals from the 1000 Genomes Project [73], or individuals sorted into superpopulations. Therefore, much of the population-specific structure is ignored, which may inflate the observed frequency differences. An adoption of a more sophisticated framework for detecting sexual antagonism, such as the one presented in [40], would be more appropriate for future studies.

Another example of genus-specific PAR1 evolution comes from the analysis of ancient human sequences. Interestingly, we find that PAR1 retained the autosomal-like phylogeny patterns of neanderthal and denisovan sequences (Figure 11) [50,74], despite the invading human PAR1 [49]. These results indicate that the invading human Y chromosome largely managed to spread in the neanderthal lineage without being linked with the human-specific PAR1 sequence. Additionally, we recover population-specific mutation patterns of archaic PAR1 sequences (Figure 13) in line with previous observations [48].

Lastly, we explore repeat content and evolution in PAR1-specific DSB regions. The mo-2 minisatellite arrays in mice [52–54] are repetitive elements involved in chromosome axes elongation and sister chromatid separation, crucial for achieving a recombination event during male meiosis. Our analysis of human sequence also implies an important recombinogenic role of repetitive elements, especially in PAR1 (Figure 14). A more detailed analysis of PAR1, using sophisticated molecular biology and microscopy methods as in [54], would be of great value for our understanding of the assurance of human PAR recombination. When analyzing DSB regions, we find patterns indicative of higher recombination rates, such as high divergence rates (Table 5) and recombination-specific substitution patterns (Figure 15). Even though these regions have been identified in human males, the observed patterns are also present in other great apes, indicating that DSB hotspots are likely conserved within the great ape lineage despite significant divergence of PRDM9-binding motifs in primates [56–58]. This observation could also indicate the existence of PRDM9-independent DSB hotspots in PAR1 across great apes, as suggested by [50]. With these results in mind, we propose that in addition to recombination map estimation and PRDM9-binding studies, repeat content analysis and comparative approaches should be included into future analyses of recombination determinants.

## Methods

### Data

Reference genomes used for alignment of the pseudoautosomal region 1 (PAR1) and autosomal telomeres, were GRCh38 for human, Clint PTRv2 for chimpanzee (University of Washington; January 2018), gorGor4 for gorilla (Wellcome Trust Sanger Institute; October 2015) Susie PABv2 for orangutan (University of Washington, 2018) and Mmul_10 for rhesus macaque (The Genome Institute at Washington University School of Medicine, 2019); https://www.ncbi.nlm.nih.gov/. We used the Progressive Cactus pipeline [59] to obtain a multiple-species alignment and ancestral sequence reconstruction. The aligned regions range from the beginning of the X chromosome to 100 kb upstream of the *XG* gene (which contains the pseudoautosomal boundary): 1-2,916,500 bp in human, 1-2,589,128 bp in chimpanzee, 1-2,486,644 in gorilla and 1-2,498,610 in macaque reference. For the PABv2 assembly, we used the unlocalized X-linked scaffold (NW_019937303.1) of length 3,293,409 which contains the orangutan PAR1 region. We only retained regions that were uniquely aligned between the sequences, which resulted in 1,066,683 base pairs of alignable sequence. We further curated the alignment by retaining only alignment blocks that are syntenic (non-inverted and non-overlapping) between the human and the three great ape species, resulting in an alignment of 620,054 sites. Finally, to obtain an alignment of putatively neutral sequences, we excluded coding regions, CpG islands, repetitive sequences and conserved regions (UCSC tracks: ncbiRefSeq, cpgIslandExtUnmasked, rmsk and phastConsElements30way). After excluding sites in and upstream of the *XG* gene, we were left with 239,799 putatively neutral PAR1 sites defined across all species and ancestral nodes. For autosomal telomeres, we chose sequences that are conserved as telomeres (3 Mb regions at the tip of an autosome) in the macaque reference [83,84]. This left us with 25 alignable autosomal telomeres with on average 872,720 putatively neutral bases for each telomere.

For human nucleotide diversity estimates we used the 30× high-coverage data of 107 individuals of the Yoruba (YRI) population from the 1000 Genomes Project [73]. The recombination map of human PAR1 was taken from [21]. The callable fraction for the human PAR1 was determined using the pilot accessibility mask of the 1000 Genomes Project, lifted-over [85] to GRCh38 coordinates.

The dataset of the eight subspecies of great apes used to estimate diversity and recombination maps was curated from four sequencing studies [69,86–88]. In total, we analyzed five subspecies of the *Pan* genus: *P. troglodytes ellioti* (N=10), *P. troglodytes schweinfurthii* (N=19), *P. troglodytes troglodytes* (N=18) and *P. troglodytes verus* (N=11) and *P. paniscus* (N=13); one subspecies of the *Gorilla* genus; *G. gorilla gorilla* (N=23); and two subspecies of the *Pongo* genus; *P. abelii* (N=11) and *P. pygmaeus* (N=15). We conducted a *de novo* read mapping and variant calling for PAR1 in all eight great ape populations as described in [71]. Recombination maps for great apes were inferred using the LDhat 2.2 program in interval mode [78], following the protocols in [76] and [89]. The callable fraction of PAR1 for great apes was determined as the number of sites covered by at least 1.5 reads per haploid genome and no more than 2×*n*×*cov* reads, where *n* is the ploidy of the region and *cov* is the mean coverage per haploid genome. We lifted-over the great ape reference sequences to GRCh38 coordinates for estimating between-species correlations of recombination and diversity (Figure 6B and D).

Estimates of autosomal nucleotide diversity (*θ*_A_) for the nine studied populations in Table 3 were taken from [69,86–88].

For the analysis of PAR1 evolution in archaic humans, we use data of the Neanderthal and Denisovan individuals sequenced to high coverage from [50,51]. The vcf coordinates of these samples were lifted-over to the GRCh38 human reference prior to analysis. The callable fraction of the archaic sequences was determined from their vcf files, which included calls of invariant sites. Coordinates of double-strand break hotspot regions in PAR1 of human males were taken from [55] and lifted to GRCh38 coordinates. For both analyses we exclude coding regions, CpG islands, repetitive sequences and conserved regions.

### Statistical analysis

All statistical analyses and plotting were conducted using the R programming software [90]. The linear model presented in Table 2 was run using the R “lm” function.

### Divergence and substitution spectra estimation

We used the program phyloFit [62] to estimate divergence rates, using the expectation maximization (option -E) and the general unrestricted single nucleotide model (--subst-mod UNREST). Additionally, we ran phyloFit using the U2S substitution model (the general unrestricted dinucleotide model with strand symmetry), to infer posterior counts and rates of different substitution types, and construct substitution spectra. To ensure convergence of the expectation maximization algorithm, we run phyloFit 10 times for each analysis with random parameter initialization (option -r) and random seed numbers (option -D). We report divergence estimates and substitution spectra for the runs with the highest likelihood values.

### Estimation of GC-biased gene conversion

Inference of the GC-biased gene conversion (gBGC) parameter *B* (4*N*_*e*_*b*, where *N*_*e*_ is the effective size of the population and *b* is the conversion parameter as defined by [36]) follows the framework in [71,91]. In short, we constructed allele frequency spectra (AFSs) with categories corresponding to GC frequency bins. For great ape samples, these bins correspond to discrete values of GC counts at a segregating site, while for the larger human population, we defined 50 equally-sized GC frequency ranges for use as AFS categories. We used only segregating sites with complete data, *i*.*e*., with defined nucleotide states for all individuals within a population. Prior to *B* inference, we omitted the two most extreme categories of AFSs as these are most likely to be affected by mutation processes and potentially bias *B* inference.

### Inference of the neighbor-joining tree of archaic sequences

We first constructed reference sequences for the Neanderthal and Denisovan individuals by converting their vcf files into fasta files using a custom python script. For polymorphic sites in the vcf, we randomly selected one of the two segregating nucleotides as the reference. To construct the neighbor-joining tree in Figure 11, we followed the protocol of [49] and used the R packages *ape* [92] and *phangorn* [93] to infer the tree and internal bootstrap support.

### Inference of repeat content

Repeat content of PAR1 and autosomal telomeres was characterized using the Tandem Repeats Finder program [75]. For all telomeres, the program was run on the unfiltered sequence (*i*.*e*., including coding regions, CpG islands, repetitive sequences and conserved regions), using the recommended parameter settings: match weight of 2, mismatch and indel penalty of 5 and 7, respectively, match and indel probability of 80 and 10, respectively, minimum alignment score of 50 and maximum period size of 2,000 bp.

## Acknowledgements

We thank Deborah Charlesworth for critical reading of the manuscript. The study was supported by grant NNF18OC0031004 from the Novo Nordisk Foundation.

## Supplement

Supplementary table 1

[telData](https://docs.google.com/spreadsheets/d/19_gjqOaBglD1fththS5SYJCXlAwRK5siTylBpDCoGEc/edit#gid=832823279)

